# Reduced expression of CorA-like metal-ion transporter homologs is associated with cation hexaammine resistance in *Plasmodium falciparum*

**DOI:** 10.64898/2026.11.23.768791

**Authors:** Manoj Yadav, Ameer Abbas, Vikas Kumar, Mangesh Bankar, Ankit Gupta

**Author notes:** Address correspondence to Ankit Gupta. Manoj Yadav, Ameer Abbas and Vikas Kumar contributed equally to this work. Author order was determined on the basis of seniority.

## Abstract

Magnesium (Mg²⁺) homeostasis is essential for parasite survival, yet its transport mechanisms in *Plasmodium falciparum* remain poorly defined. CorA-family Mg^2+^ transporters are established targets of cation hexaammines in bacteria, providing a pharmacological approach to investigate CorA-like proteins in malaria parasites. Cation hexaammines exhibited parasiticidal activities, and Co(III)Hex selection yielded a resistant parasite line with an approximately seven-fold increase in IC_50_ and a substantial growth defect. No coding mutations were detected in *Pf*MIT1, *Pf*MIT2, or *Pf*MIT3; instead, resistance was associated with reduced *Pf*MIT1 and *Pf*MIT2 transcript expression levels. Structural analyses revealed conserved CorA features, including the GMN motif and two C-terminal transmembrane helices. *Pf*MIT1 localized to both the apicoplast and mitochondrion, whereas *Pf*MIT2 and *Pf*MIT3 localized predominantly to the apicoplast. Membrane fractionation further demonstrated distinct membrane-associated states among the three proteins. Collectively, these findings implicate CorA-like Mg²⁺ transporters in parasite biology and identify *Pf*MIT proteins as candidates for antimalarial target development.

## INTRODUCTION

The pursuit of new therapeutic targets is crucial in the fight against malaria, particularly as drug resistance continues to undermine the efficacy of existing treatments. Cationic hexaammines, such as cobalt (III) hexaammine, have been reported as potent and selective inhibitors of the CorA magnesium transport system, most likely acting through competitive inhibition of Mg^2+^ uptake (1). The CorA family of Mg transporters is ubiquitous among prokaryotes, with functional homologs identified in eukaryotes. Multiple CorA homologs have been identified in eukaryotes, including five in yeast and ten in *Arabidopsis thaliana* (2, 3). These homologs are localized to distinct subcellular compartments, including the mitochondria, chloroplasts, and endoplasmic reticulum, where they play essential roles in regulating intracellular Mg²⁺ distribution and homeostasis. CorA is a dominant, constitutively expressed Mg transporter under normal conditions, whereas other transporters, e.g. MgtA and MgtB, are transcriptionally induced in response to Mg deficiency. (4). CorA family proteins form a funnel-shaped homopentamer (5). It has a large n-terminal cytoplasmic domain followed by two c-terminal transmembrane helices, which are connected by a short extracellular loop (5). The loop consists of a highly conserved Gly-Met-Asn (GMN) motif, which is essential for Mg uptake (6). The Asn (N) residue acts as a fine selectivity filter, allowing the entry of only partially hydrated Mg (7).

Magnesium (Mg) is the most abundant divalent cation among all living cells and serves as a cofactor for over 300 enzymatic reactions. In addition to its fundamental roles in cellular physiology, magnesium has also been implicated in the pathogenesis of malaria. A number of *in vitro*, animal, and clinical studies suggested a close association between Mg homeostasis and malaria infection (8–10). Serum Mg^2+^ levels have been reported to increase approximately threefold during uncomplicated acute *falciparum* malaria (11). While, a meta-analysis found significantly higher Mg^2+^ levels in patients with fatal malaria than in survivors (12). Furthermore, the Mg-deficient diet has been shown to protect mice against *Plasmodium chabaudi* infection, highlighting the importance of Mg^2+^ availability in parasite survival and disease progression (13). These observations suggest that Mg^2+^ acquisition and homeostasis are critical for *Plasmodium* biology, making Mg^2+^ transport systems attractive targets for antimalarial intervention. Despite the apparent importance of Mg^2+^ in malaria pathogenesis, relatively little is known about Mg^2+^ transport in *Plasmodium*. Three CorA homologs have been identified in *P. falciparum*, and functional studies in *P. berghei* demonstrated that these transporters are largely dispensable during blood stages (14, 15). In the present study, we investigated the parasiticidal activity of cationic hexaammines against *Plasmodium falciparum*. To explore their mechanism of action, we generated a parasite line resistant to one of the cationic hexaammines and performed molecular characterization of the resistant parasite line, which revealed downregulation of CorA homologs expression. In addition, we systematically characterized the *P. falciparum* CorA homologs and determined their subcellular localization, providing new insights into the organization of Mg^2+^ transport pathways in the malaria parasite.

## MATERIALS AND METHODS

### Homology modeling, computational and phylogenetic analysis

*P. falciparum* CorA homolog protein sequences were derived from the PlasmoDB site (https://plasmodb.org). The protein sequence alignment was performed using CLUSTALW and the ESPript3 tool. Transmembrane (TM) domains were predicted in these homologs using TMHMM2.0 (16). Homology modeling of *P. falciparum* CorA homologs was performed using MODELLER v9.24. The three-dimensional structures were generated based on the crystal structure of the CorA magnesium transporter from *Thermotoga maritima* (TmCorA; PDB ID: 3JCF) as the template. Multiple models were generated for each protein, and the model with the lowest MODELLER objective function (MolPdf) score was selected for further analysis. The stereochemical quality of the selected models was assessed by Ramachandran plot analysis using PROCHECK. To evaluate the structural accuracy of the models, root mean square deviation (RMSD) values were calculated only for the structurally aligned regions. The models showed high structural similarity to the template, with RMSD values of 0.640 Å, 0.616 Å, and 1.282 Å for *Pf*MIT1, *Pf*MIT2, and *Pf*MIT3, respectively. Phylogenetic analysis of these homologs was performed using MEGA version 12. Protein sequences were aligned using the MUSCLE algorithm implemented in MEGA12, and a phylogenetic tree was constructed using the Neighbour-Joining method with 1,000 bootstrap replicates to assess branch support.

### *In vitro* parasite culture

*P. falciparum* (3D7) parasites were cultured at 37°C under 5% O_2_, 5% CO_2_, 90% N_2_ in O^+^ human erythrocytes, RPMI 1640 (Invitrogen) supplemented with 25 mM HEPES, 50 μg/ml hypoxanthine, 0.5% Albumax (Gibco) and 0.23% NaHCO_3_. The parasite culture was synchronized using 5% sorbitol. Late-stage parasite-infected erythrocytes were enriched using the Percoll (Cytiva)-sorbitol (40% and 70%) method. The parasite pellet was enriched by lysis of the parasite culture with 0.05% saponin for 5 min on ice, followed by a spin at 6000 × g for 5 min at 4^°^C.

### Molecular cloning and transfection

Total RNA was extracted from *Plasmodium falciparum* 3D7 parasites and used for first-strand cDNA synthesis (17)*. Pf*MIT1 and *Pf*MIT2 gene fragments were PCR amplified from synthesized cDNA, whereas the *Pf*MIT3 (PF3D7_1427600) gene fragment was amplified from genomic DNA (gDNA) isolated from 3D7 parasites (18). For transient transfection, plasmid constructs were generated by cloning the gene of interest downstream of the 5′ *PfCRT* promoter in the cPARL-GFP vector, with a C-terminal GFP tag. Following validation by restriction digestion and Sanger sequencing, plasmid constructs were transfected into *Plasmodium falciparum* 3D7 parasites, and transgenic parasites were selected using 1.5 nM WR99210 (Jacobus Pharmaceuticals) (19).

### Selection of Co(III)Hex-Res parasite lines

*P. falciparum* 3D7 parasites were cultivated with 1 mM Co(III)hexaammine, resulting in microscopic clearance of parasites. Cultures were maintained under continuous drug pressure, with half of the culture volume removed and replenished with fresh erythrocytes weekly. Viable parasites capable of surviving Co(III)Hex selection were detected on day 22 by examination of Giemsa-stained blood smears. Following further expansion of the surviving population, limiting-dilution cloning was used to generate a Co(III)Hex-Res parasite line.

### Hypotonic lysis and membrane fractionation assay

Trophozoite-stage parasites were enriched by the Percoll–sorbitol method, yielding ∼ 99% infected erythrocytes and ensuring equivalent loading of synchronized parasites (20). Percoll-enriched infected erythrocytes (trophozoite stage) were lysed in ice-cold hypotonic lysis buffer (7.5 mM NaH_2_PO_4_, 1 mM EDTA, pH 7.5) containing 1 mM phenylmethylsulfonyl fluoride (PMSF) and the lysate was separated by SDS-PAGE for immunoblotting. For membrane fractionation, the hypotonic lysate was ultracentrifuged at 100,000 × g for 1 h at 4°C to obtain the soluble fraction (SF). The pellet was subsequently extracted with 100 mM Na_2_CO_3_, pH 11.0 for 30 min at 4°C and ultracentrifuged under the same conditions to separate carbonate-soluble peripheral membrane proteins (supernatant) from carbonate-insoluble integral membrane proteins (pellet). Fractions were neutralized with 1 M HCl, resolved by SDS–PAGE, transferred to nitrocellulose membranes, and immunoblotted with mouse anti-GFP (1:1000; ThermoFisher) followed by HRP-conjugated secondary antibody (1:5000; Cell Signaling Technology). Immunoreactive bands were detected using Clarity Western ECL substrate (Bio-Rad) on X-ray film.

### Immunofluorescence assay

For colocalization with the apicoplast marker *P. falciparum* histone-like protein (*Pf*HU), parasites were fixed with 4% paraformaldehyde containing 0.0075% glutaraldehyde for 30 mins at RT, and immunolabelled with rabbit anti-GFP antibody (1:250; Abcam), followed by goat anti-rabbit IgG Alexa Fluor 594 (1:100; Thermo Fisher Scientific). Additionally, for mitochondrial colocalization, parasites were stained with 50nM MitoTracker Red CMXRos (Invitrogen) for 30 mins prior to fixation as described above, followed by labelling with mouse anti-GFP antibody (1:500 dilution, Thermo Fisher Scientific) and goat anti-mouse IgG Alexa Fluor 488 (1:100; Thermo Fisher Scientific). Nuclei were counterstained with Hoechst 33342 (Thermo Fisher Scientific), and slides were mounted using ProLong Diamond antifade mountant (Invitrogen). Fluorescence images were acquired using a Zeiss confocal microscope equipped with a 64x oil-immersion objective.

### Growth rate and inhibition assays

*In vitro* growth inhibition assays were performed with synchronized ring-stage *Plasmodium falciparum* parasites (0.5% parasitemia and 2.5% hematocrit) and serial dilutions of cation hexaammines. Cultures were maintained at 37°C for 72 h, followed by addition of lysis buffer (20 mM Tris-HCl, 10 mM EDTA, 0.016% saponin, and 1.6% Triton X-100; pH 7.5) containing SYBR Green I nucleic acid gel stain (4 µL/10 mL, Invitrogen). Plates were incubated in the dark for 30 min at RT, and parasite growth was quantified by measuring fluorescence at excitation and emission wavelengths of 485 and 528 nm using a Synergy microplate reader (BioTek Instruments). Growth inhibition was calculated relative to untreated and chloroquine-treated controls, and IC_50_ values were determined by nonlinear regression using GraphPad Prism. For stage-dependent growth analysis, wild-type (3D7) and Co(III)Hex-Res parasite cultures were maintained up to 5 days and parasite growth was monitored daily using the method described above. % Growth was calculated from triplicate wells after normalization to matched cultures treated with 20 µM chloroquine.

### Quantitative real-time PCR

Expression of *PfMIT* homologs in wild-type 3D7 and Co(III)Hex-Res parasites was quantified by qRT-PCR using total RNA isolated 25 h after sorbitol synchronization with the PureLink RNA Mini Kit (Ambion). Matched late trophozoite-stage cultures were confirmed microscopically by Giemsa-stained smears. Genomic DNA was removed using the TURBO DNA-free Kit (Ambion), and first-strand cDNA was synthesized from ∼ 400 ng RNA using SuperScript III reverse transcriptase and oligo(dT) primers (Invitrogen). qRT-PCR was performed using the QuantiTect SYBR Green PCR Kit (Qiagen) on a CFX96 Real-Time PCR System (Bio-Rad) with gene-specific primers generating ∼ 121 bp amplicons (**Table S1**). Reactions were performed in triplicate and Pf07_0073 as the reference gene. Relative transcript abundance was calculated according to 2^(mean CT value of Pf07_0073 –mean CT value of the gene of interest) and expressed as arbitrary units (mean ± SEM) from independent RNA harvests.

## RESULTS

### Cation hexaammines exhibited parasiticidal activities

Cationic hexaammines, previously reported as potent and selective inhibitors of the CorA Mg^2+^ transport system, were evaluated for their parasiticidal activity on *in vitro* blood culture of *P. falciparum* 3D7 line (**Fig. 1**). Ru(III)Hex exhibited the greatest antiparasitic activity, with an IC_50_ of 22.87 ± 7.75 µM, followed by Co(III)Hex (63.80 ± 1.00 µM) and Ru(II)Hex (82.47 ± 7.35 µM) (**Table S2**). Notably, Ru(III)Hex displayed approximately 3.6-fold and 2.9-fold greater parasite killing than Co(III)Hex and Ru(II)Hex, respectively. The substantially lower IC_50_ value of Ru(III)Hex compared with Ru(II)Hex indicates that the oxidation state of ruthenium markedly influenced the biological activity of the hexaammine scaffold (**Fig. 1B-C**). For comparison, Co(III)Hex and Ru(III)Hex have previously been reported to completely inhibit the growth of *S. typhimurium* at 30 µM in Mg^2+^ deficient media (50µM), using a MM1927 strain that is solely dependent on CorA for Mg²⁺ uptake (**Table S2**) (1). The relatively higher concentrations required to inhibit *P. falciparum* may reflect differences in the experimental systems, including the substantially higher Mg^2+^ concentration in the parasite culture medium (∼ 400 µM) and the multiple biological barriers that these cationic complexes must traverse to reach their intracellular target, including the erythrocyte membrane, parasitophorous vacuole membrane, and parasite plasma membrane, etc. These factors possibly reduce the effective intracellular concentration of these compounds and thereby contribute to the higher IC_50_ values in *P. falciparum*.

**Fig 1.**
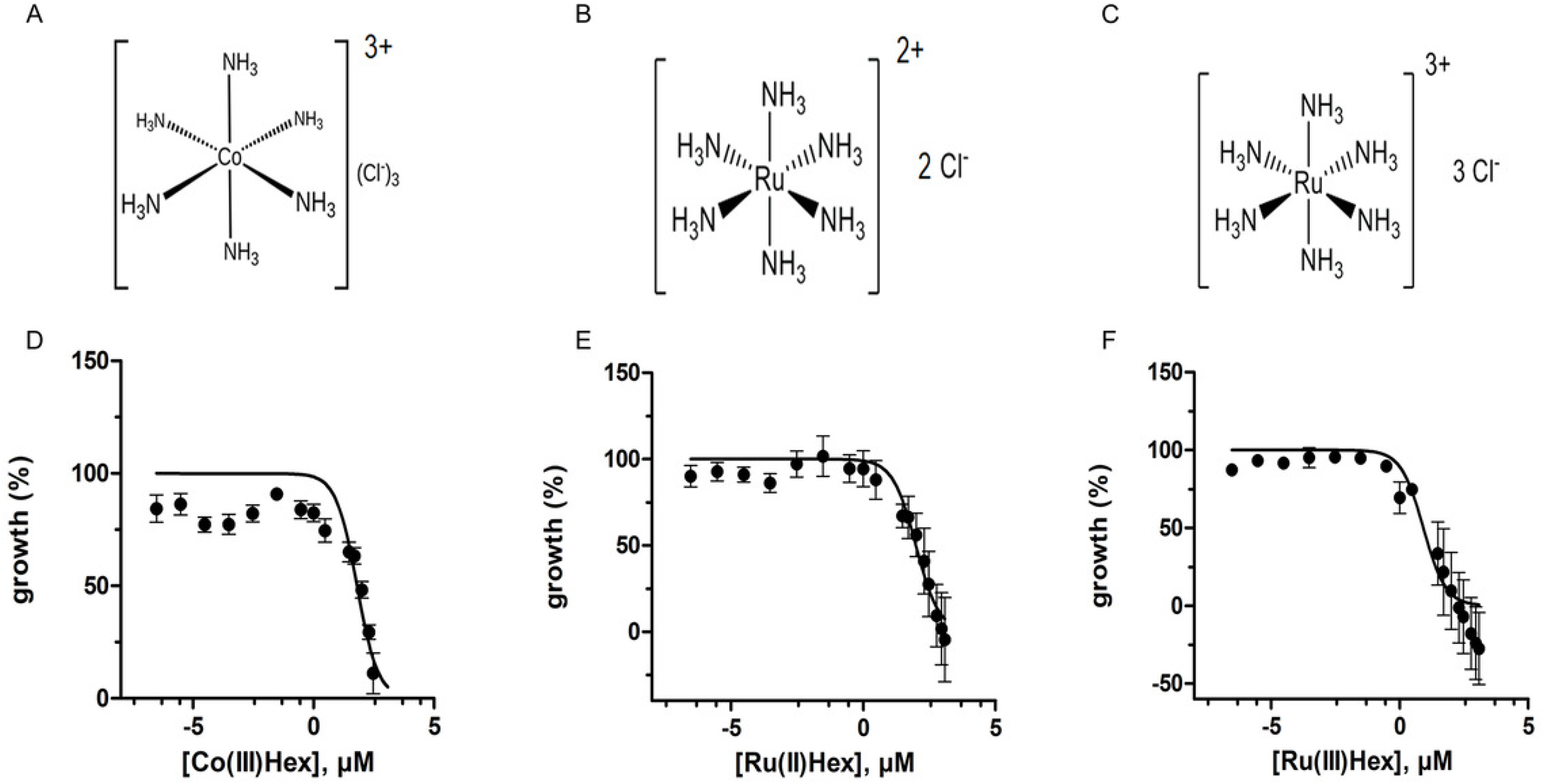
Parasiticidal activity of cation hexaammines against *P. falciparum* 3D7 parasites. Chemical structures of the cation hexaammines evaluated in this study: (A) cobalt(III)hexaammine chloride (Co(III)Hex), (B) ruthenium(II)hexaammine chloride (Ru(II)Hex), and (C) ruthenium(III)hexaammine chloride (Ru(III)Hex). Mean ± S.E.M. parasite growth at indicated concentrations of (D) Co(III)Hex, (E) Ru(II)Hex, and (F) Ru(III)Hex, respectively, determined from 3-4 independent experiments performed in triplicate with normalization to 100% for untreated control and 0% for matched cultures treated with 20 μM chloroquine for 72 h.

### Characterization of the Co(III)Hex-resistant parasite line

To gain molecular insight into cation hexaammine-mediated parasite killing, Co(III)Hex was selected to generate a resistant parasite line, based on its established structural and functional association with CorA protein-mediated Mg^2+^ transport (21). Co(III)Hex is a structural analogue of hexahydrated Mg^2+^ and has been shown by crystallographic studies to outcompete Mg^2+^ at the pore site (1, 21). We therefore used *in vitro* selection of *P. falciparum* 3D7 cultures with Co(III)hexaammine and detected outgrowth of parasites capable of surviving Co(III)Hex drug pressure after 22 days. Limiting-dilution cloning of the surviving population yielded the Co(III)Hex-Res line, which exhibited substantially reduced sensitivity to Co(III)Hex compared with the parental 3D7 line (**Fig. 2A**). Co(III)Hex selection produced an IC_50_ of 484.40±55.54 µM for the Co(III)Hex-Res line, compared with 77.18±14 µM for the parental 3D7 line, corresponding to ∼ 7-fold decrease in Co(III)Hex sensitivity in the Co(III)Hex-Res line (P < 0.05; **Fig. 2A-B**). Additionally, Co(III)Hex-Res parasites showed ∼60% reduced growth compared with parental wild-type 3D7 parasites (**Fig. 2C**).

**Fig 2.**
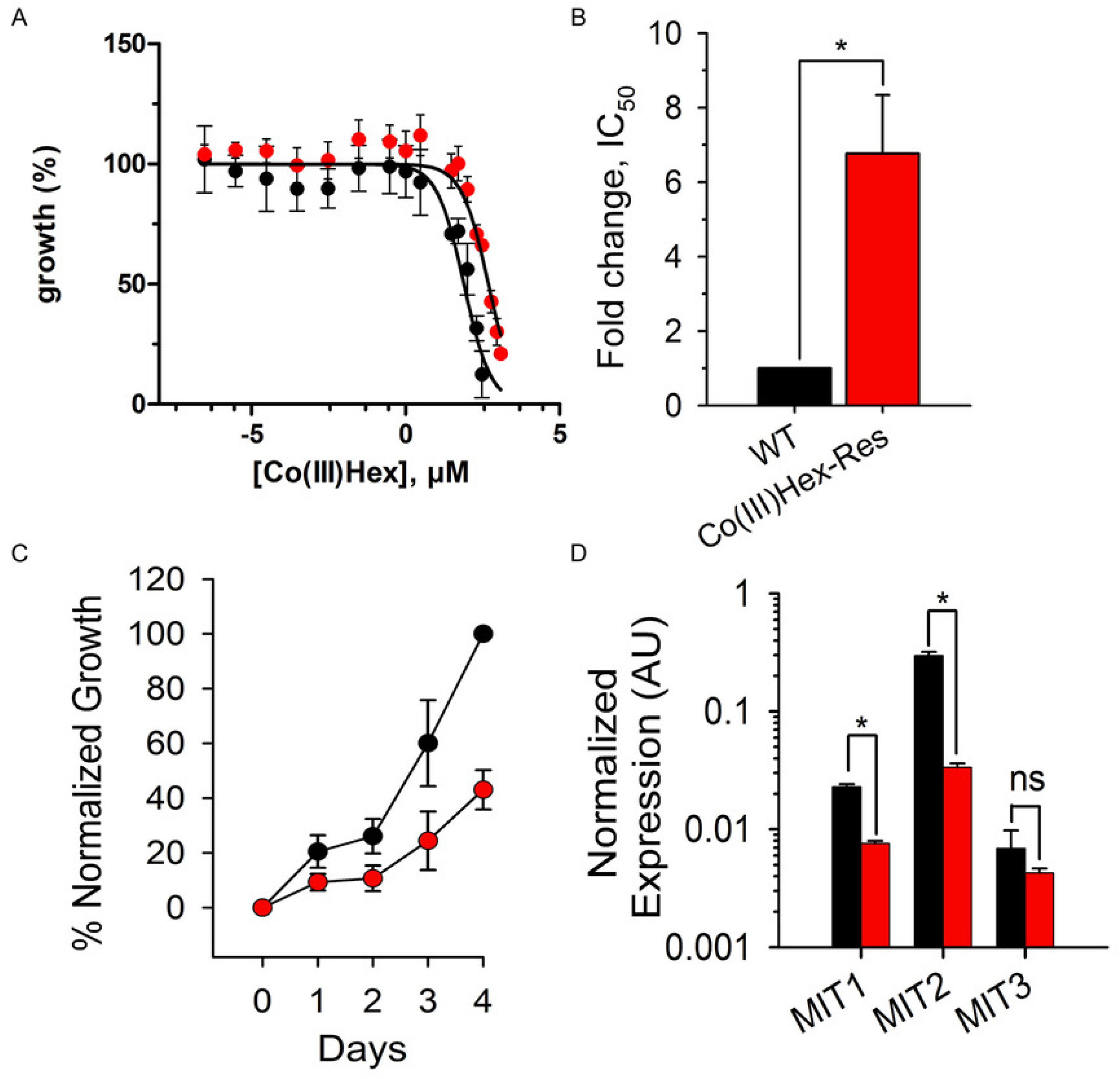
Selection and phenotypic characterization of Co(III)Hex-Res parasites. (A) Mean ± S.E.M. parasite growth of wild-type (WT; black circles) and Co(III)Hex-resistant (Co(III)Hex-Res; red circles) parasites at the indicated Co(III)Hex concentrations, determined from 3–4 independent experiments, each performed in triplicate. Parasite growth was normalized to 100% for untreated controls and 0% for matched cultures treated with 20 μM chloroquine for 72 h. (B) Fold change in IC_50_ values for WT and Co(III)Hex-Res parasites, calculated from the dose–response curves shown in panel A. Bars represent mean ± S.E.M. (n=3-4); Asterisk, p = 0.02. (C) Mean ± S.E.M. normalized parasite growth of WT (black circles) and Co(III)Hex-Res (red circles) parasites over 4 days (n=3). (D) Mean ± S.E.M. normalized expression of indicated *Pf*MIT1, *Pf*MIT2, and *Pf*MIT3 genes in wild-type (black bar) and Co(III)Hex-Res (red bar) parasites, calculated according to the 2^−ΔCt^ method using PF07_0073 as the endogenous reference gene (internal loading control); n = 4 independent biological replicates. Asterisks indicate statistically significant differences (*P* < 0.05, Student’s *t*-test); ns, not significant.

To further examine the molecular basis of Co(III)Hex-resistance, the full-length *Pf*MIT1, *Pf*MIT2, and *Pf*MIT3 genes were PCR-amplified and sequenced in the Co(III)Hex-Res and parental 3D7 parasites (data not shown). We selected these genes based on their similarity to bacterial CorA proteins and published structural data identifying residues involved in Co(III)Hex interaction with the CorA pore region (21). No mutations were identified in any of the three *Pf*MIT genes in the Co(III)Hex-Res line relative to parental 3D7 parasites, suggesting that the reduced sensitivity to Co(III)Hex was not associated with coding-sequence mutations in these CorA homologs. We next examined the transcript expression levels of the three *PfMIT* genes. Expression of *PfMIT1* and *PfMIT2* was significantly reduced in Co(III)Hex-Res parasites compared with the parental 3D7 parasites, whereas *PfMIT3* expression did not differ significantly between these two parasite lines (**Fig. 2D**). Thus, despite the absence of detectable coding-sequence mutations in *Pf*MIT1, *Pf*MIT2, and *Pf*MIT3, Co(III)Hex resistance was associated with selective downregulation of *PfMIT1* and *PfMIT2*, but not *PfMIT3*. These findings suggest that reduced expression of parasite CorA homolog proteins, particularly *Pf*MIT1 and *Pf*MIT2, is associated with reduced sensitivity to Co(III)Hex in *P. falciparum*, in the absence of detectable mutations in their coding sequences. However, whether these transcriptional changes directly contribute to resistance or represent a downstream response to Co(III)Hex exposure remains to be determined.

### *Pf*MIT homolog proteins exhibit conserved structural features of CorA-family Mg^2+^ transporters

Phylogenetic analysis revealed that *Pf*MIT1 was positioned between the α-proteobacterial and plant/green-algae CorA proteins, suggesting a mixed mitochondrial- and plastid-associated evolutionary signal (**Fig. 3A**). In contrast, *Pf*MIT2 and *Pf*MIT3 were more closely associated with apicomplexan and plant/green-algal CorA proteins and lacked a clear association with the α-proteobacterial clade, indicating a predominantly plastid-associated evolutionary signal (**Fig. 3A**). Collectively, these findings suggest that *Pf*MIT1, *Pf*MIT2, and *Pf*MIT3 have followed distinct evolutionary trajectories, with *Pf*MIT1 potentially reflecting contributions from both mitochondrial and plastid lineages, whereas *Pf*MIT2 and *Pf*MIT3 are more closely associated with the plastid-related lineage. Consistent with the phylogenetic analyses, *in silico* targeting predictions further supported the dual targeting of *Pf*MIT1 to both the apicoplast and mitochondria, with PlasmoAP predicting no signal peptide and a strong targeting peptide (-/++) and MitoProt II predicting a high probability of mitochondrial targeting (98.29%) (**Table S3**).

**Fig 3.**
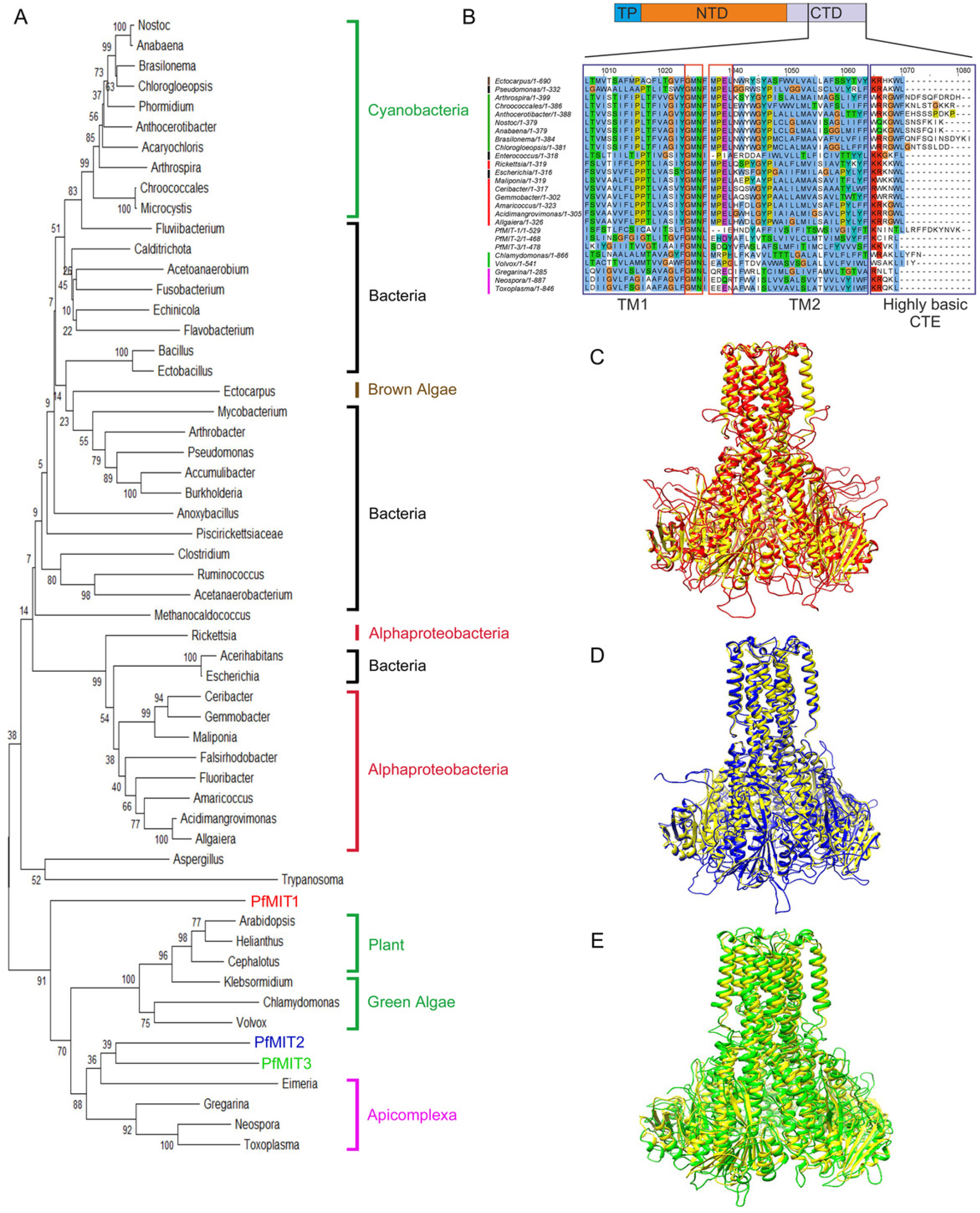
Phylogenetic and structural analyses of *Plasmodium falciparum* CorA homologs. (A) Phylogenetic tree showing the evolutionary relationships of *Pf*CorA homologs (*Pf*MIT1, red; *Pf*MIT2, blue; *Pf*MIT3, green) identified across bacteria, cyanobacteria, brown algae, green algae, plants, apicomplexans, and other eukaryotes using the Neighbor-Joining method. Major taxonomic groups are indicated by coloured brackets on the right. The evolutionary distances were computed using the JTT matrix-based method and are in units of amino acid substitutions per site. The optimal tree with the sum of branch lengths = 46.578 is shown. Bootstrap support values (%) from 1,000 replicates are indicated at branch nodes. (B) Multiple sequence alignment (ClustalW) of CorA proteins highlighting conserved structural features. Predicted protein architecture consists of an N-terminal domain (NTD), two transmembrane helices (TM1 and TM2), and a highly basic C-terminal extension (CTE). (C– E) *Pf*MIT1 (red), *Pf*MIT2 (blue), *Pf*MIT3 (green) were modeled on the crystal structure of *Thermotoga maritima* CorA (yellow) protein using MODELLER v9.24.

Based on multiple sequence alignment, *Pf*MIT1, *Pf*MIT2, and *Pf*MIT3 exhibited 10.96%, 12.39%, and 12.34% protein sequence identity and similarity of 20.98%, 26.28%, and 23.43%, respectively, with *Thermotoga maritima* CorA (*Tm*CorA) (**Table S4**). Notably, *Pf*MIT2 showed the highest protein sequence identity and similarity, whereas *Pf*MIT1 exhibited the lowest, consistent with the evolutionary relationships observed in the phylogenetic analysis. Despite the relatively low overall sequence identity and similarity, the C-terminal domain (CTD) of the *Pf*MIT homologs retained the characteristic conserved GMN motif, two predicted transmembrane domains (TM1 and TM2) and a highly basic C-terminal region, which are conserved structural features of CorA family proteins (**Fig. 3B**). Notably, *Pf*MIT1 carries a distinct elongated C-terminal extension (CTE) that closely resembles those of cyanobacterial and green-algal CorA proteins but differs from the CTEs observed in other apicomplexan homologs, including *Pf*MIT2 and *Pf*MIT3, as well as in bacterial CorA proteins (**Fig. 3B**). In addition to these conserved features, sequence alignment revealed that the highly conserved MPEL motif, located within the conserved interhelical loop between TM1 and TM2, was absent in *Pf*MIT1, *Pf*MIT2, and *Pf*MIT3 (**Fig. 3B**). The MPEL motif was likewise absent in several bacterial and green-algal CorA homologs, indicating that its absence is not unique to apicomplexan MITs and may reflect lineage-specific variation within the CorA protein family.

CorA-family Mg^2+^ transporters form funnel-shaped homopentameric channel assemblies in which each monomer contains two C-terminal transmembrane helices (TM1 and TM2) that contribute to the central ion-conducting pore, together with a large N-terminal cytosolic domain involved in Mg^2+^ sensing and regulation through a negative-feedback mechanism (22). Homology modelling based on *Tm*CorA revealed that *Pf*MIT1, *Pf*MIT2, and *Pf*MIT3 retain the characteristic CorA-family architecture and are predicted to form homopentameric assemblies, comprising a membrane-spanning C-terminal domain with two transmembrane helices (TM1 and TM2) per monomer and a larger N-terminal cytosolic domain (**Fig. 3D–E**). Despite their sequence divergence and distinct evolutionary relationships, all three *Pf*MIT homologs retained a broadly conserved structural organization that may contribute to their interaction with cationic hexaammines.

### Differential Subcellular Localization of *Pf*MIT1, *Pf*MIT2, and *Pf*MIT3 to the Apicoplast and Mitochondrion

To experimentally assess the subcellular localization of the three *Pf*MIT homologs, transgenic parasite lines overexpressing *Pf*MIT1, *Pf*MIT2, and *Pf*MIT3 proteins with a C-terminal GFP tag were generated through transient transfection using the *CpARL-GFP* plasmid (**Fig. 4A, D, G**). Here, anti-Hu was used as an apicoplast marker (23). Immunofluorescence analysis showed that the *Pf*MIT1-GFP signal overlapped with both anti-Hu and MitoTracker signals, suggesting that *Pf*MIT1 localizes to both the apicoplast and mitochondrion (**Fig. 4B i–ii**). Western blot analysis confirmed the expression of GFP-tagged *Pf*MIT1 at ∼90 kDa, with an additional band corresponding to autocleaved GFP detected in lysates from the transgenic parasites but absent in the wild-type control (**Fig. 4C**). An overlap was observed between the *Pf*MIT2-GFP signal and anti-Hu, with no overlap detected with MitoTracker, consistent with the predicted apicoplast localization of the *Pf*MIT2 protein (**Fig. 4E i-ii**). Western blot analysis confirmed the expression of GFP-tagged *Pf*MIT2, with the expected band at ∼82 kDa, and another band of autocleaved GFP in the transgenic parasite lysate but not in the wild-type control (**Fig. 4F**). Similarly, the *Pf*MIT3-GFP signal overlapped with the anti-Hu signal but showed no overlap with the MitoTracker signal, suggesting that *Pf*MIT3 localizes exclusively to the apicoplast and not to the mitochondrion **(Fig. 4H i-ii)**. Western blot analysis revealed a band of ∼ 84 kDa corresponding to GFP-tagged *Pf*MIT3 in the transgenic parasite lysate, with no evidence of GFP autocleavage (**Fig. 4I**). This band was absent in the wild-type control. Anti-actin was used to verify equal protein loading.

**Fig 4.**
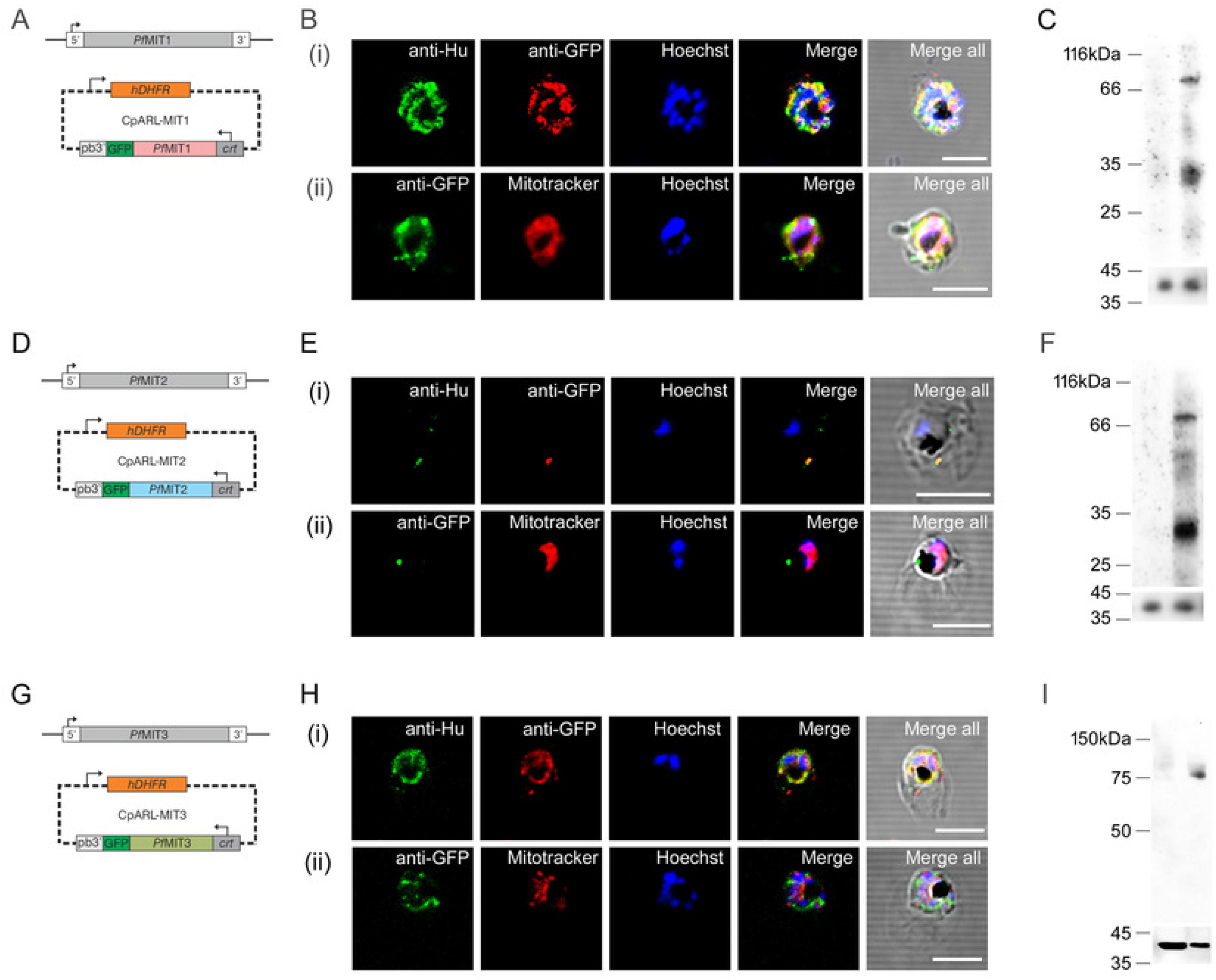
Subcellular localization of *Pf*MIT1, *Pf*MIT2, and *Pf*MIT3 in *Plasmodium falciparum*. (A) *Pf*MIT1 was transiently overexpressed as a C-terminal GFP fusion under the control of the PfCRT promoter. (B) Overlap of *Pf*MIT1-GFP signal with (i) apicoplast marker (anti-Hu) and (ii) mitotracker suggests its dual localization to both apicoplast and mitochondrion. (C) Western blot analysis of the *Pf*MIT1-GFP transfectant showing expression of the fusion protein (∼90.5 kDa) and an autocleaved GFP product (lane 2), with no corresponding signal detected in the wild-type control (lane 1). β-Actin served as the loading control in both lanes. (D) Transient transfection with cPARL-MIT2 resulted in overexpression of *Pf*MIT2-GFP. (E) Colocalization of the GFP signal with (i) anti-Hu and (ii) absence of overlap between the GFP and MitoTracker signals indicate that *Pf*MIT2 is targeted to the apicoplast but not to the mitochondrion. (F) Western blot analysis showed no signal in the wild-type control (lane 1), whereas expression of the *Pf*MIT2-GFP fusion protein was observed at ∼82 kDa, along with an autocleaved GFP product (lane 2). β-Actin was used as the loading control to confirm equal protein loading. (G) Schematic depicting transient transfection strategy for overexpression of *Pf*MIT3-GFP. (H) Colocalization of the GFP signal with (i) anti-Hu and (ii) the absence of overlap with the MitoTracker signal indicate that *Pf*MIT3-GFP is exclusively targeted to the apicoplast, not to the mitochondrion. (I) Western blotting confirmed expression of the *Pf*MIT3-GFP fusion protein (∼84 kDa) in the transfectant (lane 2), whereas no corresponding band was detected in the wild-type control (lane 1). β-Actin was used as the loading control in both lanes. Scale bars, 5 μm.

### Two-state behaviour and predicted Membrane Topology of *Pf*MIT1, *Pf*MIT2, and ***Pf*MIT3**

Hypotonic lysis released *Pf*MIT1 and *Pf*MIT2 into the soluble fraction (SF) with an additional band corresponding to autocleaved GFP detected in both cases (**Fig. 5A-B**, first lane in each panel). Following Na_2_CO_3_ treatment, *Pf*MIT1 was predominantly detected in the carbonate-insoluble pellet (CI, third lane) fraction, whereas only a small amount of *Pf*MIT2 was detected in this fraction, with no signal observed in the carbonate-soluble (CS, second lane) fraction (**Fig. 5A–B**). Thus, *Pf*MIT1 and *Pf*MIT2 appear to exist in two distinct states: a soluble form and an integral membrane-associated form. In contrast, *Pf*MIT3 was not detected in the soluble fraction following hypotonic lysis (**Fig. 5C**, first lane). Upon Na_2_CO_3_ treatment, *Pf*MIT3 was detected predominantly in the carbonate-soluble (CS) fraction, with a comparatively weaker signal in the carbonate-insoluble pellet (CI) fraction (**Fig. 5C**, second and third lanes, respectively). These findings suggest that *Pf*MIT3 exists predominantly as a peripheral membrane-associated protein, with a smaller fraction present as an integral membrane protein.

**Fig 5.**
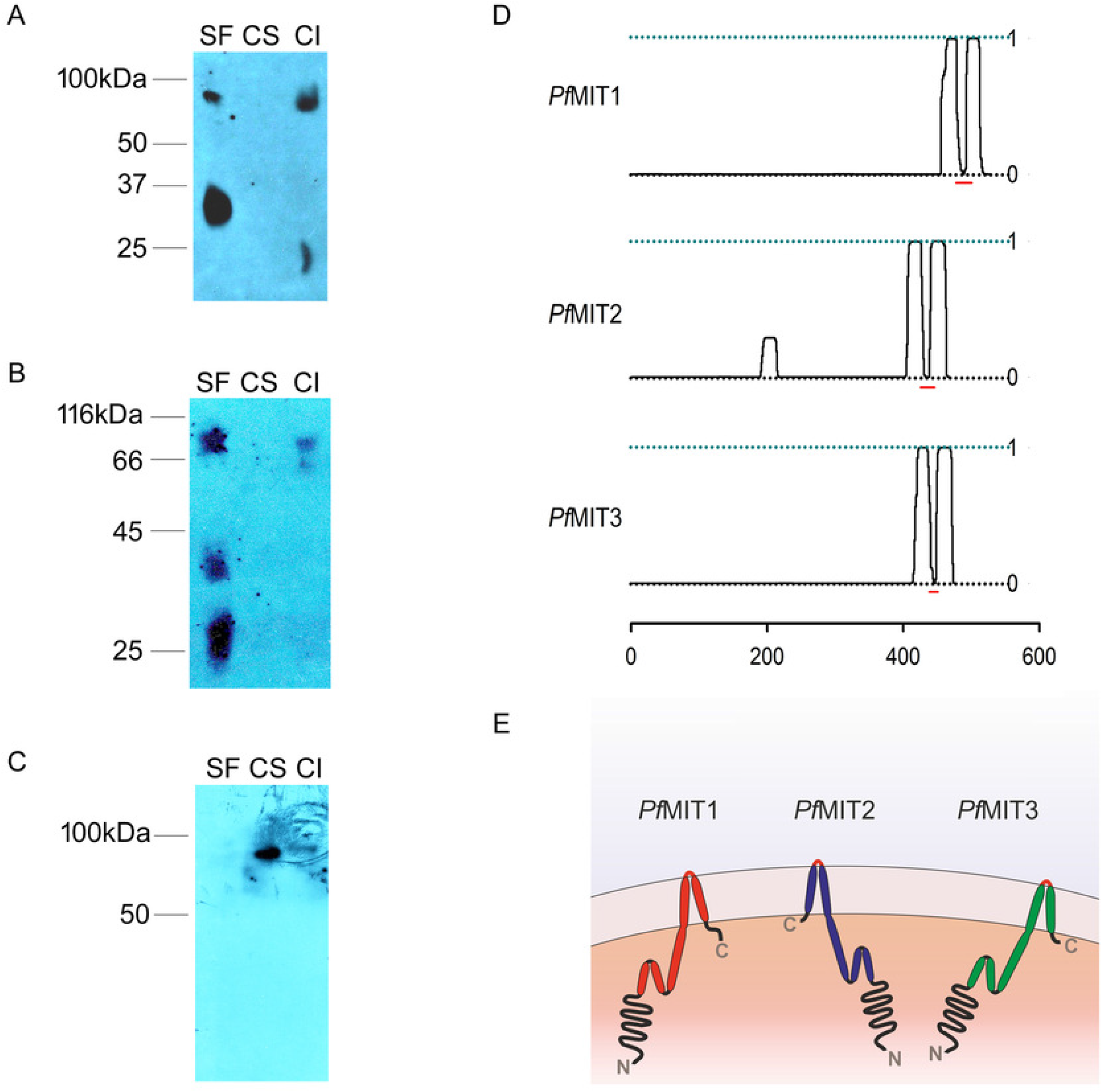
Membrane association and topology of *Pf*MIT1, *Pf*MIT2, and *Pf*MIT3. (A-C) An equal number of Percoll-enriched parasites from each transgenic line were subjected to hypotonic lysis followed by alkaline treatment, yielding soluble (SF), carbonate-extractable (CS), and integral membrane (CI) fractions. (D) Posterior probability plots showing the predicted transmembrane topology of each MIT homolog, as determined using the Phobius algorithm. Solid black lines denote predicted TM regions, whereas the red line indicates the intervening membrane-exposed region between TM1 and TM2. Residue positions are shown relative to the unprocessed N-terminus of each *Pf*MIT homolog. (E) Schematic depicting a proposed membrane topology shared among the three *Pf*MIT homologs. Transmembrane helices are shown as colored cylinders (*Pf*MIT1, red; *Pf*MIT2, blue; *Pf*MIT3, green), with the N- and C-termini indicated. *Pf*MIT1 contains an extended C-terminal region enriched in basic amino acids compared with *Pf*MIT2 and *Pf*MIT3.

The membrane fractionation assay results were consistent with computational analyses predicting two hydrophobic regions (TM1 and TM2) toward the C-terminus on both sides of this site (**Fig. 2D**, red dashed lines), which are predicted to be exposed outside in each *Pf*MIT homolog. Notably, *Pf*MIT2 also contained an additional short hydrophobic region within its N-terminal portion. Based on these computational predictions, all three *Pf*MIT homologs were modelled as membrane proteins containing two C-terminal transmembrane helices, with relatively long N-terminal and shorter C-terminal regions oriented toward the cytosol (**Fig. 2E**). Interestingly, *Pf*MIT1 possessed a comparatively long, basic amino acid-rich C-terminal extension, unlike *Pf*MIT2 or *Pf*MIT3. Collectively, the membrane fractionation and topology analyses support the membrane-associated nature of *Pf*MIT homologs and further corroborate their classification as divergent CorA-family Mg^2+^ transporters.

## DISCUSSION

A number of studies underscore the importance of Mg^2+^ acquisition and homeostasis in *Plasmodium* survival and disease progression; yet, the mechanisms governing Mg^2+^ transport in *P. falciparum* remain poorly defined (8–10, 13). Here, we identify three CorA-like proteins: *Pf*MIT1, *Pf*MIT2 and *Pf*MIT3, which share conserved structural features including the “GMN” motif characteristic of the CorA/Mrs2/Alr1 superfamily, and exhibit distinct organellar localization (24). Importantly, the inhibition of parasite growth by cationic hexaammines and the emergence of a Co(III)Hex-Res parasite line associated with reduced *Pf*MIT1 and *Pf*MIT2 transcript levels provide pharmacological and genetic evidence linking these proteins to Mg^2+^ homeostasis and parasite biology.

Cationic hexaammines have been reported to selectively inhibit CorA-mediated Mg^2+^ uptake by mimicking hydrated divalent cations and interacting with the channel pore (1, 21). Thus, the antiparasitic activity of Co(III)Hex, Ru(II)Hex, and Ru(III)Hex against *P. falciparum* suggests that a CorA-like Mg^2+^ transport process may contribute to parasite survival. The ∼3-fold difference between the Ru(II)Hex and Ru(III)Hex further indicates that the chemical properties of the metal centre strongly influence antiparasitic activity, consistent with their known effects on CorA interaction. However, the higher IC_50_ values observed against *P. falciparum* compared with *S. typhimurium*, should not necessarily be interpreted as reduced target affinity. Unlike bacterial CorA proteins, the parasite targets are localized to subcellular organelles and require compounds to traverse multiple membrane barriers, while the higher extracellular Mg²⁺ concentration in parasite cultures may further reduce their effective intracellular concentration and increase the apparent IC_50_. Notably, the free ionic Mg^2+^ concentration in human serum (∼8.5-11 mg/L) is substantially lower than that in RPMI medium (48.84 mg/L), suggesting that the higher Mg^2+^ concentration under *in vitro* culture conditions may contribute to the reduced sensitivity of *P. falciparum* to cationic hexaammines.

The generation of a Co(III)Hex-Res parasite line provides genetic evidence linking Co(III)Hex sensitivity to CorA-like Mg^2+^ transporters in *P. falciparum*. Resistance was associated with an ∼7-fold increase in the Co(III)Hex IC_50_, indicating adaptation to sustained drug pressure. Notably, the Co(III)Hex-Res parasites exhibited a significant reduction in growth rate compared with the parental line, suggesting a substantial fitness cost associated with resistance. Sequencing revealed no coding mutations in *the Pf*MIT1*, Pf*MIT2*, or Pf*MIT3 genes; instead, the resistant line showed significantly reduced transcript levels of *Pf*MIT1 *and Pf*MIT2, suggesting that altered expression of these CorA-like transporters may contribute to Co(III)Hex resistance. This pattern is consistent with reduced target abundance as a potential mechanism of resistance, analogous to metronidazole resistance in *Gardnerella vaginalis* (25), and is consistent with observations in bacterial CorA, where reduced CorA-mediated Mg^2+^ uptake can alter susceptibility to divalent-cation-associated toxicity (26).

Nevertheless, these findings establish an association rather than causality. Reduced transcript levels may result from altered promoter/terminator activity, changes in the activity of linked transcriptional regulators modulating protein abundance, or an adaptive mechanism that decreases the cellular abundance of drug-sensitive transporters. Alternatively, these changes could reflect a secondary response to altered Mg^2+^ homeostasis or broader physiological adaptations induced by drug selection. Direct genetic manipulation and complementation of *Pf*MIT1 and *Pf*MIT2, coupled with quantitative assessment of Mg^2+^ transport, will be required to distinguish these possibilities.

The crystal structure of bacterial CorA revealed a funnel-shaped homopentameric assembly in which two transmembrane helices (TM1 and TM2) from each subunit form the central ion-conducting pathway (5, 21). The GMN motif contributes to ion selectivity, whereas the cytoplasmic domain senses intracellular Mg^2+^ to regulate channel gating (24). Consistent with this architecture, the *PfMIT* homologs retain key CorA features, including the GMN motif, two C-terminal transmembrane helices, a basic C-terminal region, and a predicted pentameric organization, supporting their classification as CorA-like proteins despite substantial sequence divergence. Highly basic CTE is a characteristic feature of CorA proteins, where positively charged residues contribute to membrane anchoring and, upon oligomerization, form a basic ring at the cytosolic membrane interface that participates in channel gating.

Notably, *Pf*MIT1 has a longer, lysine-rich C-terminal extension than *Pf*MIT2 and *Pf*MIT3, which may confer distinct membrane interactions, oligomeric stability, or regulatory properties; however, these possibilities require experimental validation.

*Pf*MIT1 is evolutionarily distinct and exhibits dual localization to the apicoplast and mitochondrion, in contrast to *Pf*MIT2 and *Pf*MIT3, which are exclusively targeted to the apicoplast. These findings are consistent with the phylogenetic and computational predictions and suggest functional diversification among the *Pf*MIT homologs. The presence of three *Pf*MIT homologs, together with a recently reported apicoplast-localized protein showing structural similarity to CorA-family metal transporters, raises the possibility that these proteins may form distinct homopentameric transporters associated with the individual membrane layers of the four-membrane apicoplast envelope, or alternatively, assemble into heteropentameric complexes to facilitate metal transport across the apicoplast membranes (27). However, these localization patterns should be interpreted cautiously, as GFP tagging and protein overexpression may influence protein targeting; therefore, further validation using endogenous tagging, organelle fractionation, or immunoelectron microscopy would provide stronger evidence for the same. *Pf*MIT1 and *Pf*MIT2 were detected in both soluble and carbonate-insoluble fractions, whereas *Pf*MIT3 was recovered in both carbonate-soluble and -insoluble fractions, broadly consistent with their predicted transmembrane architecture. The differential fractionation may reflect differences in membrane association, stability, or interactions with other parasite proteins.

## CONCLUSIONS

Cationic hexaammines exhibited parasiticidal activity against *P. falciparum*, and selection with Co(III)Hex generated a resistant parasite line with reduced *Pf*MIT1 and *Pf*MIT2 transcript levels and a substantial fitness cost. The three *Pf*MIT homologs retained conserved CorA-like structural features but exhibited distinct evolutionary relationships, organellar localization, and membrane-associated properties, suggesting functional diversification. Collectively, these findings support a role for CorA-like proteins in parasite biology and identify *Pf*MIT homologs as candidates for further investigation as components of Mg^2+^ homeostasis and potential antimalarial targets.

## ACKNOWLEDGEMENTS

We acknowledge all volunteers and the Blood Bank at AIIMS Raebareli for providing fresh blood used for parasite culture. We also acknowledge Dr Saman Habib (CSIR-CDRI, Lucknow) for providing WR99210 and Ms Rima for helping us with confocal microscopy imaging. The work was funded by Start-up Research Grant (SRG/2019/001574) and Core Research Grant (CRG/2022/001219) received from the Science and Engineering Research Board (SERB), India.

## Supplementary Material

Tables S1-S4: (i) Details of primers used in qRT-PCR. (ii) IC_50_ values for cation hexaammines in *P. falciparum* (3D7) and *S. typhimurium*. (iii) *In silico* prediction of organellar targeting in *Pf*MIT homologs (iv) Sequence identity/similarity of *P. falciparum* MIT homolog proteins.

## NOTES

The authors declare no competing interests. The Institutional Ethics Committee (PGI/IEC/2019/001739) approved the study protocol for work involving human blood in parasite culture, and the Institutional Biosafety Committee (2026/IBC/2019) approved it for generating transgenic parasite lines.

